# Extensive Secondary Contact Among Three Glacial Lineages of Arctic Char (*Salvelinus alpinus*) in Newfoundland and Labrador

**DOI:** 10.1101/394742

**Authors:** Sarah J. Salisbury, Gregory R. McCracken, Donald Keefe, Robert Perry, Daniel E. Ruzzante

**Author notes:** Corresponding author: Sarah J. Salisbury Address: Dept. of Biology, Dalhousie University, 1355 Oxford St., Halifax, NS, B3H 4R2, Canada.

## Abstract

We sequenced a portion of the D-loop region in over 1000 Arctic char (*Salvelinus alpinus*) samples from 67 locations across Newfoundland and Labrador to assess the extent of secondary contact among the Arctic, Atlantic, and Acadian glacial lineages in Newfoundland and Labrador. Within Labrador, the Arctic and Atlantic lineages were widespread. Two locations (one landlocked and one with access to the sea) also contained individuals of the Acadian lineage, constituting the first record of this lineage in Labrador. Atlantic and Acadian lineage individuals were found in both eastern and western Newfoundland. Multiple sampling locations in Newfoundland and Labrador contained fish of two or more different glacial lineages, implying their introgression. Glacial lineage did not appear to dictate contemporary genetic divergence between the pale and dark morph of char present in Gander Lake, Newfoundland. Both were predominately of the Atlantic lineage, suggesting the potential for their divergence in sympatry. This study reveals Newfoundland and Labrador to be a unique junction of three glacial lineages which have likely hybridized extensively in this region.

## INTRODUCTION

Glaciation events are a significant driver of evolution, physically isolating species into separate glacial refugia which can undergo allopatric divergence for thousands of years (Hewitt 2000, 2004; Fraser et al. 2012). During allopatry, populations may experience differential selection and drift resulting in the formation of genetically distinct glacial lineages (Hewitt 2003, Ruzzante et al. 2008, Moore et al. 2015). Retreating glaciers allowed glacial lineages to expand into previously inaccessible environments, sometimes resulting in their secondary contact (Hewitt 2000, Swenson and Howard 2005, Soltis et al. 2006). Upon secondary contact, glacial lineages may freely hybridize, leading to complete genomic introgression (Hewitt 1988). However, if significant genetic differences have accrued between glacial lineages they may have achieved reproductive isolation prior to secondary contact or secondary contact may reinforce reproductive isolation due to selection against hybrids (Hewitt 1988, Noor 1999, Schluter 2001). Gene flow between glacial lineages in secondary contact may lie at some intermediate level between these two extremes (i.e. full hybridization and reproductive isolation) depending on the adaptive quality of each gene (Hewitt 1988). The amount of genetic divergence accumulated between glacial lineages and the degree of erosion of this divergence in secondary contact zones can significantly influence the contemporary genetic structure of a species (Bernatchez and Wilson 1998, Hewitt 2000, 2004). Since the degree of introgression is dictated by the adaptive and neutral genetic divergence between lineages, these areas of secondary contact and hybridization offer natural experiments for the study of the factors driving speciation (Hewitt 1988).

Arctic char (*Salvelinus alpinus*) is one such species whose evolutionary history has been shaped by the allopatric divergence of glacial lineages during the Pleistocene (Brunner et al. 2001, Moore et al. 2015). Five glacial lineages of Arctic char have been described based on mtDNA: Arctic, Atlantic, Acadian, Beringian, Siberian (Brunner et al. 2001, Moore et al. 2015). Evidence for secondary contact has been observed between the Beringian and Arctic lineages in Russia and western North America (Brunner et al. 2001, Moore et al. 2015, Esin et al. 2017, Oleinik et al. 2017) and between the Arctic and Atlantic lineages in Nunavut and Labrador, Canada and between the Atlantic and Acadian lineages in Newfoundland, Canada (Wilson et al. 1996, Brunner et al. 2001, Moore et al. 2015, Salisbury et al. 2017). However, our knowledge of these secondary contacts is at a coarse spatial scale, particularly in Atlantic Canada. Between Wilson et al. (1996), Brunner (2001) and Moore et al. (2015) fewer than 100 individuals from 7 locations were assessed for glacial lineage in Atlantic Canada. Salisbury et al. (2017) assessed 96 individuals at a further 3 locations. Only 3 of these 10 locations (Fraser River (Wilson et al. 1996), Ikadlivik River (Moore et al. 2015) and Ramah (Salisbury et al. 2017)) were found to contain fish of multiple glacial lineages. These locations represent the only locations across the entire species’ range where Arctic char of different glacial lineages have been observed to cooccur. Therefore the extent of secondary contact and introgression among glacial lineages remain largely unexplored in Newfoundland and Labrador.

The Laurentide Ice Sheet covered this region during the Pleistocene (Bryson et al. 1969). It retreated fully from Newfoundland between 13 000 and 9 000 years BP (Bryson et al. 1969, Dyke 2004, Shaw et al. 2006) and from Labrador between 9 000 – 7 500 years BP (Bryson et al. 1969, Jansson 2003, Occhietti et al. 2011). The vast quantities of freshwater draining from the retreating glaciers into the Atlantic Ocean allowed anadromous Arctic char to extensively colonize Newfoundland and Labrador (Power 2002). Contemporarily, this species inhabits lakes in Newfoundland and Labrador (some of which have lost their access to the sea) and where char remain in freshwater year-round (Scott and Crossman 1998). Labrador also contains anadromous populations in sea-accessible locations (Bernatchez et al. 1998).

These anadromous Arctic char populations are economically significant in Labrador where they form the basis of a commercial, recreational and subsistence fishery (Coady and Best 1976, LeDrew 1980, DFO 2001, Dempson et al. 2008). Historic allopatry may be an important underlying influence on the genetic structure of the Arctic char populations contributing to these fisheries if char of different glacial lineages contribute to the fishery but remain reproductively isolated. Such cryptic population structure could necessitate the development of independent management plans for different glacial lineages. It is currently unknown which glacial lineages contribute to the Labrador fishery, but this knowledge is potentially critical for its management.

The relative contribution of different glacial lineages to anadromous versus landlocked populations also remains largely unknown in this region. Landlocked and anadromous populations might have been founded by different lineages, for example, if one was better adapted to a particular environment or life history. Alternatively, landlocked populations could have been founded only by lineages that were present before access to these lakes was lost. Therefore investigation of which lineages are present in these two types of populations may give an indication of the timing of colonization of different glacial lineages (Moore et al. 2015).

Newfoundland and Labrador are also known to contain char exhibiting within-lake genetic structure. Salisbury et al. (2017) found evidence of genetic structure in two landlocked lakes and one sea-accessible lake in Labrador. Morphological evidence suggested that the genetic structure in the sea-accessible lake corresponded to resident and anadromous morphs. In both cases, the observed genetic structure was uncorrelated with glacial lineage. Glacial lineage has also been suggested as the origin of the substantial genetic divergence observed between a pale and a dark morph documented for Gander Lake in Newfoundland (Gomez-Uchida et al. 2008).

We therefore sought to investigate the secondary contact and potential hybridization of the Arctic, Atlantic and Acadian glacial lineages across Newfoundland and Labrador at a fine spatial scale. We hypothesized that if Labrador was colonized from the north by the Arctic lineage and from the south by the Atlantic lineage, then the Arctic lineage would be more prevalent in northern populations of char than the Atlantic lineage. We looked for locations containing both the Atlantic and Acadian lineage in Newfoundland and both the Atlantic and Arctic lineage in Labrador to support the hybridization between each of these pairs of lineages in these regions. In Labrador we sought to compare the lineages present in landlocked versus sea-accessible locations. If either the Arctic or Atlantic lineage colonized Labrador after the sea-accessibility of contemporarily landlocked lakes had been lost, then the late-colonizing lineage should only be found in anadromous populations. Finally, within Gander Lake we investigated the glacial lineage of the pale and dark morphs for evidence that their divergence is the result of their founding by different glacial lineages. To test these hypotheses we employed mtDNA to identify the glacial lineage of hundreds of fish across Newfoundland and Labrador.

## METHODS

### Sampling

Tissue samples (N = 1329) were collected between 2000 and 2015 from Newfoundland and Labrador. Landlocked and sea-accessible Labrador locations were distributed among 10 drainages (fjords or bays). The samples from 3 sites in Labrador (Ramah (R01), WP132 (S03), WP133 (S04) were used previously in Salisbury et al. (2017). Collections from western Newfoundland originate from five landlocked lakes in the Upper Humber River (see Gomez-Uchida et al 2009, Gomez-Uchida et al 2013). Collections from eastern Newfoundland originate from two locations containing only freshwater residents: Gander lake (including samples of the pale and dark morphs described for this lake (Gomez-Uchida et al 2008)), and Wing Pond.

Samples from Labrador were collected using electrofishing in the rivers (sea-accessible sites) and variably-sized standardized nylon monofilament gillnets (1.27 cm to 8.89 cm diagonal) at the landlocked and sea-accessible lake sites. Samples were collected from anadromous char populations in the Okak and Voisey regions as well as from the Fraser River, Anaktalik River, and Tikkoatokak River. These populations contribute to the three stock complexes of the commercial Labrador char fishery (Okak, Voisey, Nain) (LeDrew 1980, DFO 2001). Gander Lake was sampled using Lundgren multimesh gillnets (Hammar and Filipsson, 1985) (bar length from 0.625 cm to 7.5 cm) (see Gomez-Uchida et al. 2008 for more details). The Upper Humber River was sampled using fyke nets and eletrofishing (see Gomez-Uchida et al. 2009 for more details). Wing Pond was sampled with gillnets. Fish were weighed, measured for fork length (FL) in mm, and assessed for sex and maturity. Tissue samples (fin or gill) were obtained and immediately stored in 95% ethanol; alternatively, some fin clip samples were stored dry. All samples were collected in collaboration with the Department of Environment and Conservation for Newfoundland and Labrador and/or Parks Canada and in accordance with Dalhousie University’s Animal Ethics Guidelines.

### DNA Extraction, Amplification and Genotyping

Tissue samples were digested at 55°C for approximately eight hours using Proteinase K (Bio Basic Inc., Markham, ON, Canada). DNA was then extracted using a Multiprobe II plus liquid handling system (Perkin Elmer, Waltham, MA, USA) using a glassmilk protocol modified from Elphinstone et al. (2003).

The left domain region of the mitochondrial control region was amplified and sequenced following Moore et al. (2015). In brief, the primers *Tpro2* (Brunner et al. 2001) and *SalpcrR* (Power et al. 2009) were used to amplify the entire control region using the thermocycler program and PCR reaction outlined in (Brown Gladden et al. 1995). A shorter fragment was amplified using *Char3* instead of *Tpro2* for a minority of samples which had poor quality as determined from visual inspection of a 1% agarose gel. For all samples, a total of ∼500 bp of the left domain was sequenced using *Char3* (Power et al. 2009) at MacrogenUSA (Rockville, MD). Each unique haplotype detected was validated by resequencing a representative sample for each haplotype using *Tpro2*. All sequences were manually trimmed and validated for accuracy.

### Analyses

Our sequences were aligned along with a reference haplotype set (including control region haplotypes verified by Moore et al. (2015) and Salisbury et al. (2017)), and the control region sequences for three other salmonid species present in the study region brook trout (*Salvelinus fontinalis*), lake trout (*Salvelinus namaycush*), and Atlantic salmon (*Salmo salar*) (for accession numbers see Table S1, Supporting Information). Sequences were aligned using the Geneious alignment and default parameters in GENEIOUS (10.0.9, Auckland, NZ, www.geneious.com). Sequences verified as Arctic char were ascribed to the reference haplotype(s) for which they had 0 basepair differences. Non-char sequences were ascribed to the brook trout, lake trout or Atlantic salmon haplotype to which they had the minimum number of basepair differences.

A representative forward sequence for each unique haplotype (i.e. those sequences which contained one or more basepair differences from those haplotypes verified by Moore et al. (2015)) was aligned with its reverse complement (sequenced with *Tpro2*) using a pairwise Geneious alignment and default parameters to create a consensus sequence. The consensus sequences for these unique haplotypes were then aligned with the reference haplotype set using a Geneious alignment. A gap penalty of 7 was used and all other parameters were kept at default values. A tree was constructed based on this alignment using the PhyML (Guindon and Gascuel 2003) plugin in GENEIOUS to compare the phylogenetic relationships among these unique consensus sequences with those haplotypes verified by Moore et al. (2015) and those of two outgroup species (brook trout and lake trout). We used the Nearest Neighbour Interchange topology search algorithm, the HKY85+I+G model, and calculated 1000 bootstraps for each node following Moore et al. (2015).

A haplotype map based on all unique haplotypes found in this study along with all haplotypes verified by Moore et al. (2015) was created using PopArt version 1.7 (Leigh and Bryant, 2015). Haplotypes were trimmed to 501 bp, the length for which all haplotypes had no missing basepairs, since PopArt masks missing basepairs. This meant that haplotype ATL04 and a unique haplotype ATL31 were indistinguishable in this analysis since the SNP differentiating these haplotypes lies outside of this 501 bp region. The haplotype map was created using a Median-Joining network (Bandelt et al. 1999) with an Epsilon value of 0.

A spatial analysis of molecular variance (SAMOVA 2.0) (Dupanloup et al. 2002) was employed to detect groups of sampling locations whose F_CT_ were maximally differentiated based on mtDNA sequences. All sequences were aligned using Geneious alignment and default parameters and trimmed to 512 bp to include all relevant SNPs differentiating haplotypes. Sampling locations with fewer than 10 sequences were excluded from the analysis to minimize the probability of biased groupings due to small sampling size. F_CT_ values were estimated using a simulated annealing optimization process for K = 2 – 10 groups for all sampling locations and for only the Labrador sampling locations and for K = 2 - 4 groups for only the Newfoundland sampling locations. For each K-value, molecular distance was calculated using Tamura and Nei distance between all sampling locations and between only those sampling locations connected using a Delaunay network (Delaunay 1934) based on the latitude and longitude of each sampling location. The use of a Delaunay network limits groupings to geographically proximate sampling locations. Simulations were run for 10 000 steps from 100 initial configurations using a missing data value of 1 (such that the entire 512 bp was included in the analysis).

Linear regressions between latitude and the number of lineages present in each location as well as binomial logistic regressions of latitude on the presence or absence (coded as 1 and 0, respectively) of each of the relevant glacial lineages were conducted using R (R Core Team, 2013) for all locations, only Labrador locations, and only Newfoundland locations.

## RESULTS

### Species distribution

Samples from 59 locations in Labrador (of which 43 are sea-accessible and 16 are landlocked (Anderson 1985)) and 8 locations in Newfoundland (all containing lacustrine residents) for a total of 67 locations overall were successfully sequenced (Table 1). Five locations in Labrador were excluded from analyses due to poor sequence quality (N = 109 individuals). A further 20 individuals from across the remaining 67 locations were excluded from analyses due to poor sequence quality. A total of N = 1296 individuals were successfully sequenced across all locations in Newfoundland and Labrador (Table 1). Of these, 1133 had haplotypes consistent with the Arctic char species. The remaining 163 individuals were identified as brook trout, lake trout, or Atlantic salmon. All locations contained at least one Arctic char haplotype except G06 which only contained Atlantic salmon. In the remaining locations we sampled between 1 - 48 Arctic char (median = 18.5), which made up between 2% - 100% of the haplotypes in each sample.

**Table 1.**
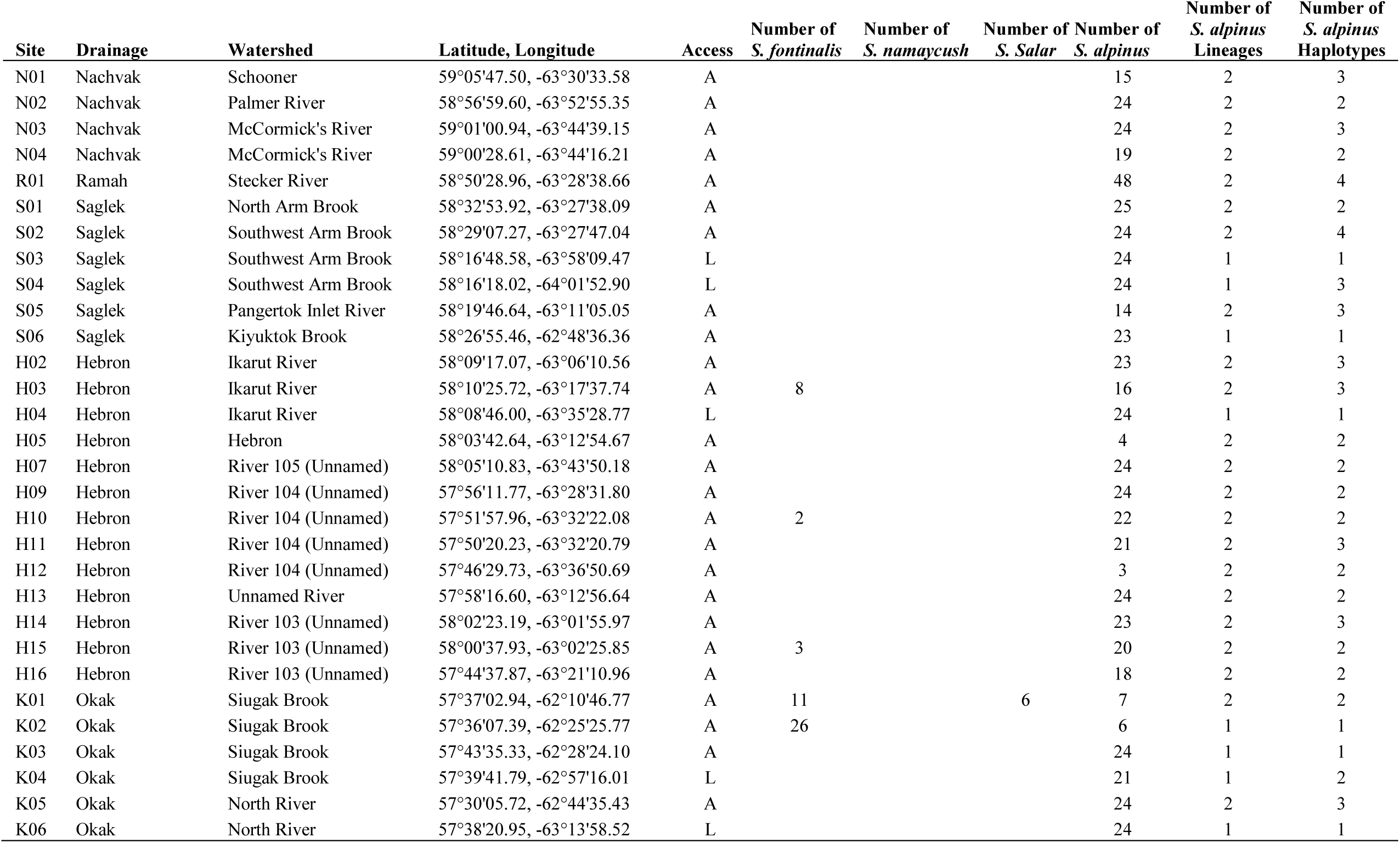
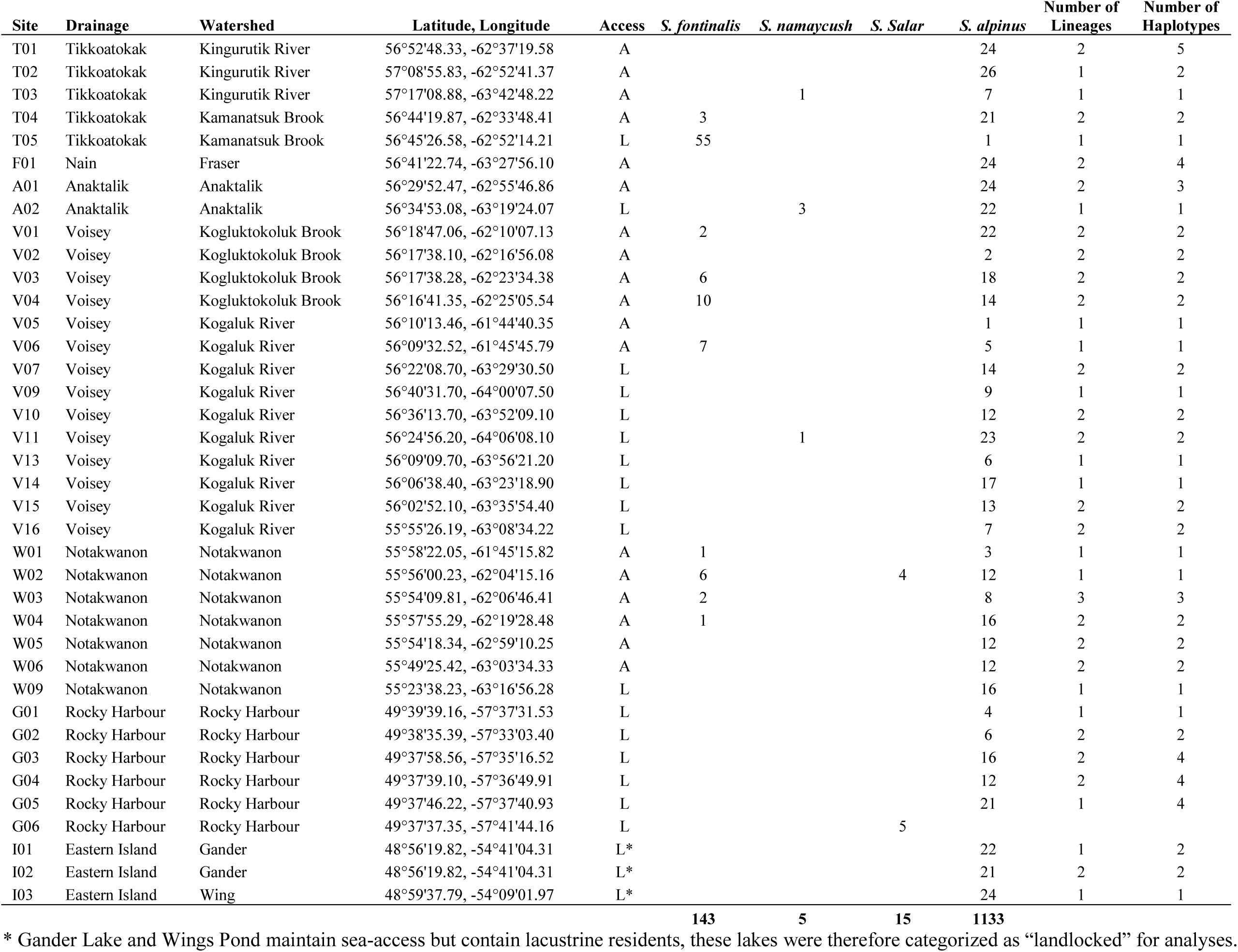
Number of Arctic char (*Salvelinus alpinus*) samples, glacial lineages and haplotypes as well as number of Brook trout (*Salvelinus fontinalis*), Lake trout (*Salvelinus namaycush*), and Atlantic salmon (*Salmo salar*) samples verified by mtDNA sequencing at sampling locations across Newfoundland and Labrador. Accessibility of locations (A for sea-accessible, L for landlocked).

### Glacial lineage distribution

The Arctic and Atlantic glacial lineages were ubiquitous across Labrador and both lineages were present in all 10 drainages (Fig.1a). The Arctic and Atlantic lineages were detected in 49 and 48 locations, respectively, and they co-occurred in 39 locations. The Acadian lineage was detected in only two sampling locations in Labrador. In one landlocked location (A02, Fig.1a), all char samples were of the Acadian lineage. The second location was sea-accessible (W03, Fig.1a) and it contained one individual of the Acadian lineage among 1 Arctic lineage and 6 Atlantic lineage individuals.

**Fig. 1.**
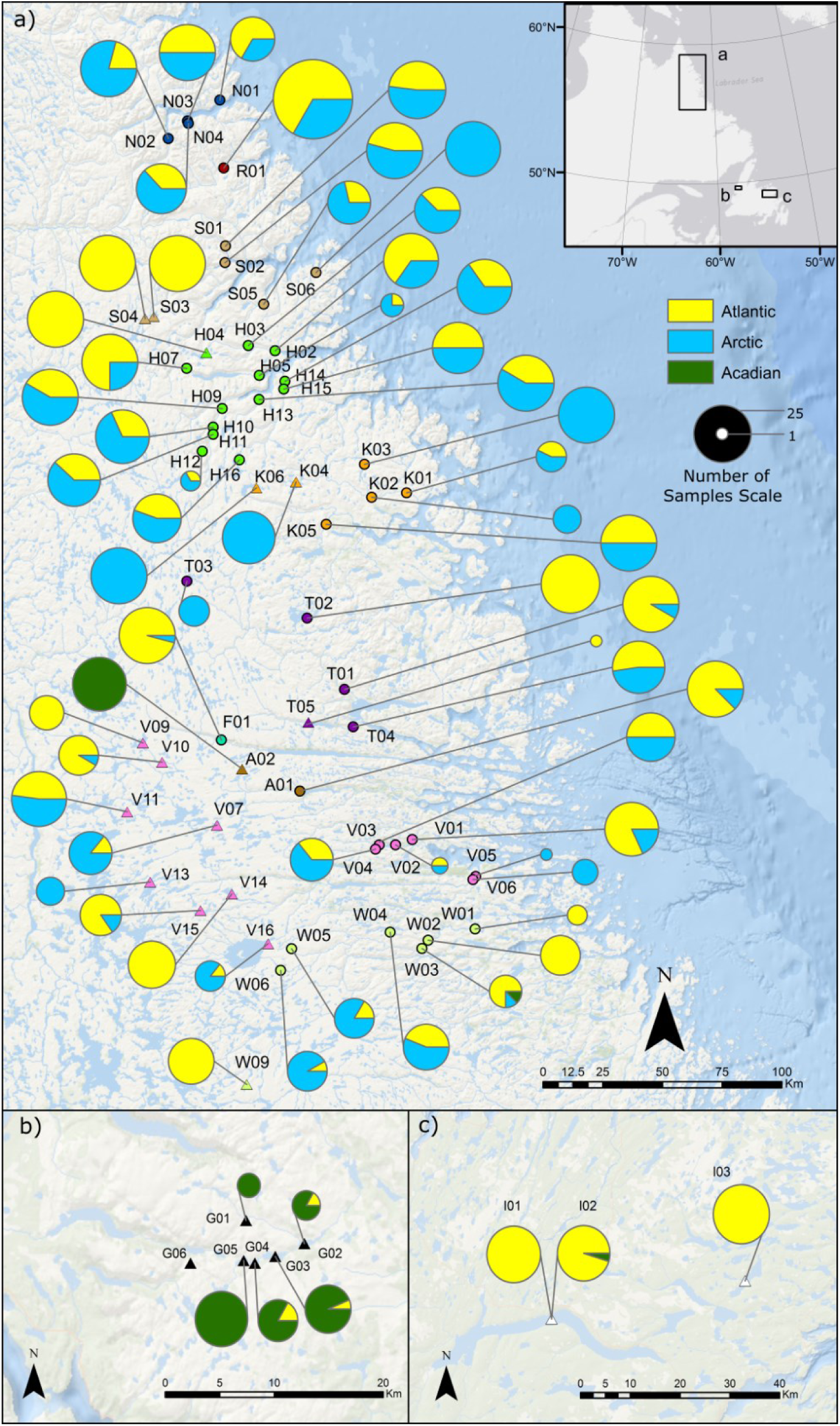
Map of sampling locations for Arctic char (*Salvelinus alpinus*) in a) Labrador and the b) west and c) east coasts of Newfoundland. Sea-accessible sites are denoted by circles, landlocked sites are denoted by triangles. Sites of the same colour are in the same drainage. Pie charts indicate the number of samples of the Acadian, Atlantic or Arctic lineage observed at a given site and are scaled by sample size. Map created using ArcGIS (ESRI).

Only the Atlantic and Acadian lineages were detected in Newfoundland (Fig.1b,c). The Atlantic lineage was detected in 3/5 locations in western Newfoundland and both locations in eastern Newfoundland. The Acadian lineage was detected in all 5 locations of western Newfoundland but only a single Acadian lineage individual was detected in a pale morph char from Gander Lake in eastern Newfoundland.

The extensive overlap in the distributions of these three lineages in Labrador is suggested by a lack of correlation between latitude and the number of lineages present in each sampling location across all locations (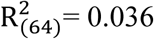, p ≥ 0.12), in only Labrador sites (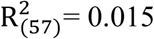 p ≥ 0.36), and in only Newfoundland sites (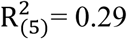, p ≥ 0.21). Binomial logistic regressions of latitude on the presence or absence (coded as 1 and 0, respectively) of the Arctic and Atlantic lineages in each Labrador sampling location were not significant (p ≥ 0.699, p ≥ 0.145, respectively). Similarly, there was no significant relationship between latitude and the presence of the Atlantic and Acadian lineages in Newfoundland sites (p ≥ 0.38, p ≥ 0.350, respectively). Across all sampling locations, the presence of the Atlantic lineage was unrelated with latitude (p ≥ 0.435). However, across Newfoundland and Labrador the probability of Arctic lineage presence increased with latitude (p ≤ 6.46 × 10^-3^). Similarly, the presence of the Acadian lineage was inversely related to latitude (p ≤ 1.08 × 10^-4^).

### Haplotype Distribution

The most common haplotype within a lineage coincided with the most common haplotypes reported in Moore et al. (2015) (haplotypes at each location: see Table S2). A total of 86% of Atlantic lineage individuals exhibited haplotype ATL01, while 78% of Acadian lineage individuals exhibited haplotype ACD9. All Labrador samples of the Acadian lineage had this haplotype. Lastly, over 99% of Arctic lineage individuals exhibited haplotypes ARC19 or ARC24. These two haplotypes were distinguished by a single SNP outside of the region sequenced using the *Char3* primer. However, the reverse complement of 29 samples from 28 locations and 9 drainages with either the ARC19 or ARC24 haplotype was sequenced using *Tpro2* and all were found to have the haplotype ARC19. Therefore, unambiguously ARC19 sequences were grouped with sequences that could be either ARC19 or ARC24 when counting the number of haplotypes present in a given site. The ARC19, ATL01, and ACD9 haplotypes were found across the modern distributions of the Arctic, Atlantic, and Acadian lineages respectively by Moore et al. (2015).

Other detected haplotypes which had been previously described by Moore et al. (2015) and Salisbury et al. (2017) include ACD11, ARC20, ARC22, ATL19, ATL23, ATL24 and ATL25. A single sample had the ATL19 haplotype, a dark morph char from Gander. This was also the only Atlantic haplotype other than ATL01 detected in Newfoundland. The ATL23, ATL24 and ATL25 haplotypes were only observed in S03 and S04 as described in Salisbury et al. (2017) except for 1 individual with ATL23 found in S02. This individual may have been washed downstream from the immediately upstream landlocked S03 and S04. This is supported by its identification as a putative migrant from S03 based on GENECLASS2 (Piry et al. 2004) results as reported in Salisbury et al. (2017).

Five samples from 3 landlocked sites in the Voisey drainage had shortened sequences that prevented their differentiation between ATL01 and ATL04. Since these sites also contained individuals unambiguously identified as ATL01, these shortened sequences were considered to be ATL01 when counting the number of haplotypes present in these lakes. The ATL04 haplotype was also found in 12 sea-accessible sampling locations across 7 drainages in Labrador. The reverse complement of 9 of these samples from 6 drainages was sequenced using *Tpro2* and all were found to contain a consistent SNP in the consensus sequence, differentiating this haplotype from ATL04. One of these samples was from R01, previously mistakenly identified as ATL04 in Salisbury et al. (2017). Given the consistency of this SNP across samples from multiple drainages, sampling locations, and studies, we denoted this as a new haplotype ATL31. We considered all ATL04 haplotypes and verified ATL31 haplotypes to be a single haplotype when counting the number of haplotypes present in the 12 sea-accessible sampling locations where these haplotypes were observed.

Including ATL31 there were 8 haplotypes not previously identified by Moore et al. (2015) or Salisbury et al. (2017) (Fig.2). All new haplotypes were one base pair different from another haplotype verified by Moore et al. (2015) within their assigned lineage (Fig.3). These include three Acadian haplotypes (ACD12, ACD13, ACD14) only observed in western Newfoundland. Four new Atlantic haplotypes were identified (ATL26, ATL28, ATL29, ATL31). ATL26 was found in only one individual in A01. ATL28 was found in three individuals, one in F01, one in T01, and one in T02. ATL29 was found in one individual in N01. Only one new Arctic haplotype, ARC35, was observed in a single individual in T01. All new haplotypes except ATL31 were found to be at least 1 base pair different from the top hit when compared with the NCBI nr/nt database using the Megablast algorithm. ATL31 was found to have 100% identity with an Arctic char sample (Accession Number: KY122252) collected from Lake Sitasjaure, Sweden (Oleinik et al. 2017).

**Fig. 2.**
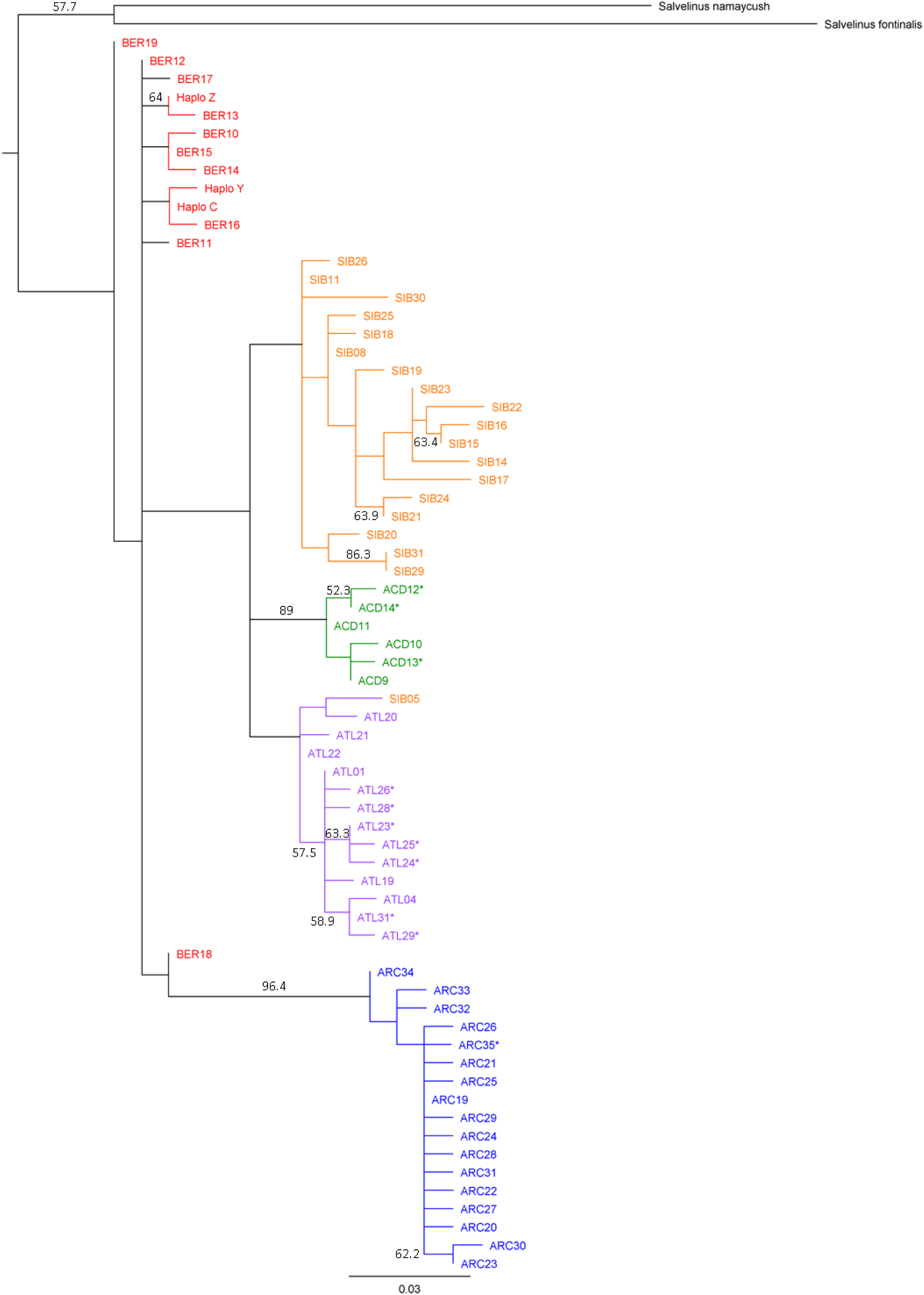
Maximum likelihood phylogenetic tree of Arctic char (*Salvelinus alpinus*) haplotypes of the mtDNA control region. Tree was generated using PhyML (Guindon and Gascuel 2003) with 1000 bootstrap replicates. Those bootstrap values greater than 50% are shown on the tree. Haplotypes are colour-coordinated by lineage as designated in Moore et al. (2015): blue - Arctic, red - Bering, orange - Siberia, purple - Atlantic, green, - Acadian. New haplotypes identified in this study and Salisbury et al. 2017 are starred.

**Fig. 3.**
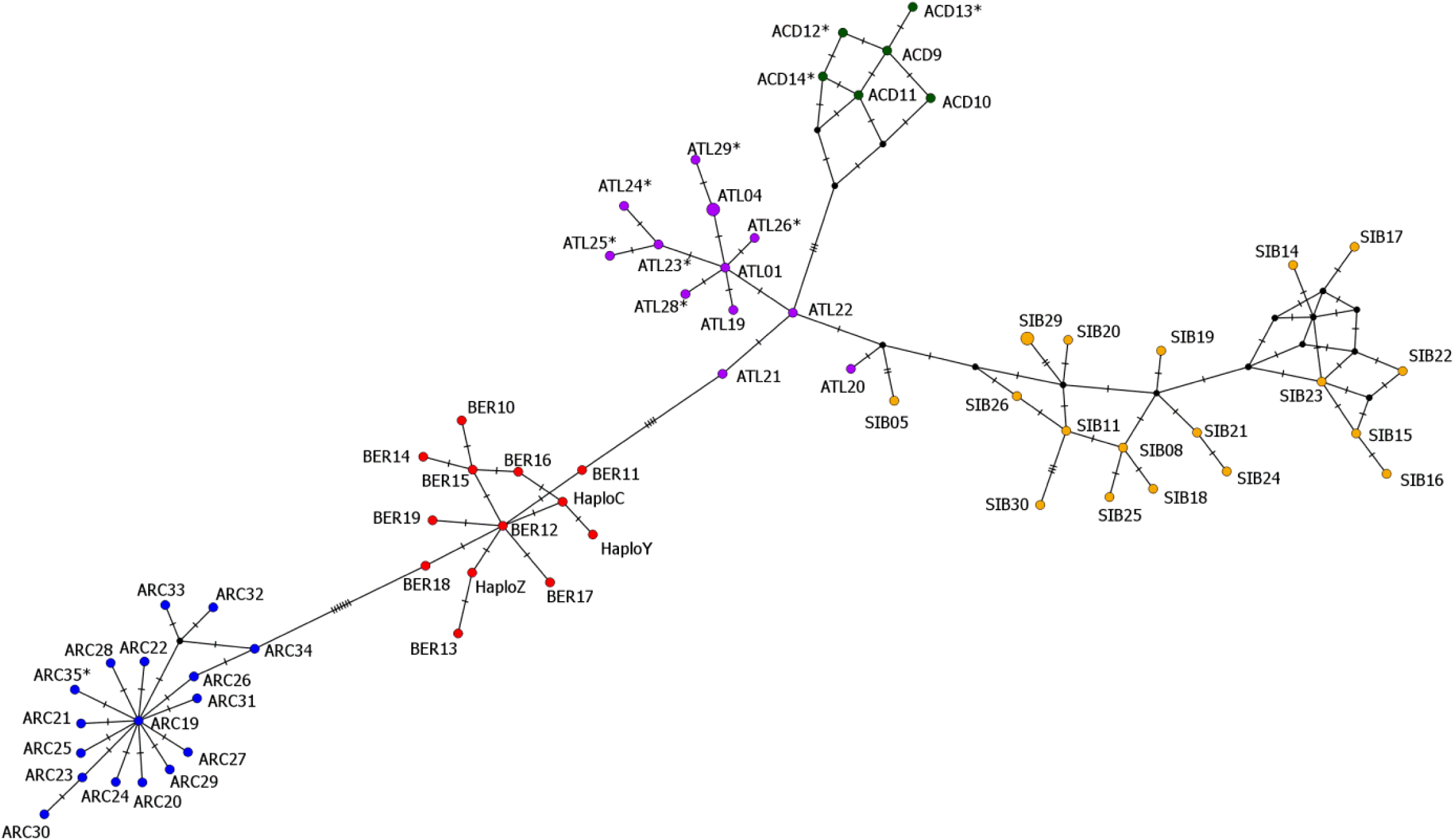
Haplotype map of Arctic char (*Salvelinus alpinus*) haplotypes created with PopArt version1.7 (Leigh and Bryant, 2015) using a Median-Joining network (Bandelt et al. 1999) and an Epsilon value of 0. New haplotypes identified in this study and Salisbury et al. 2017 are starred.

### Landlocked versus Sea-accessible Sampling Locations

The average number of lineages observed in anadromous sites was higher than in landlocked sites (average of 1.8 lineages for anadromous sites versus 1.3 lineages for landlocked sites, T_(42)_ = 3.78, p ≤ 0.001). Similarly, the average number of haplotypes observed in anadromous sites was higher than in landlocked sites (average of 2.3 haplotypes for anadromous sites versus 1.8 haplotypes for landlocked sites, T_(45)_ = 1.96, p ≤ 0.056). (Note: the Gander lake morphs (I01/I02) and Wing Pond were grouped with the landlocked lakes despite both lakes having access to the sea because their char are lacustrine residents.) This effect was even more extreme when considering only Labrador sites where the corresponding average numbers were 2.3 and 1.5 haplotypes in anadromous and landlocked sites, respectively (T_(40)_ = 3.56, p ≤ 0.001). Anadromous sites in Labrador also had an average of 1.8 lineages per site, significantly more than the 1.3 lineages observed in landlocked sites (T_(26)_ = 3.63, p ≤ 0.0012).

## SAMOVA

SAMOVA results were similar across all samples and when considering Newfoundland and Labrador sampling locations separately. Results were also similar with and without the use of a Delaunay network to take into account geographic proximity of locations. For brevity, we report only the SAMOVA results when considering all sampling locations and a Delaunay network (for results of all other SAMOVA analyses see Fig.S1, S2).

When considering all sampling locations, F_CT_ was maximized for K = 6 (Fig. 4). However, the difference in F_CT_ between K = 6 and K = 5 was small (i.e., 0.13) and a plot of F_CT_ versus K revealed that F_CT_ peaked at K = 5 (Fig. S1a). Given this small difference in F_CT_ we report the more parsimonious results of K = 5. The first group contained 26 populations across Labrador with approximately equal proportions of Arctic and Atlantic lineage individuals. The second group contained only Arctic lineage individuals and comprised 4 locations in Labrador, three in the Okak and one in the Saglek drainage. The third group comprised only one Labrador site, A02, which contained only Acadian lineage individuals. The fourth group contained 16 populations, 14 in Labrador and 2 in eastern Newfoundland. This group comprised largely Atlantic lineage individuals. The fifth group contained the three landlocked lakes in western Newfoundland.

**Fig. 4.**
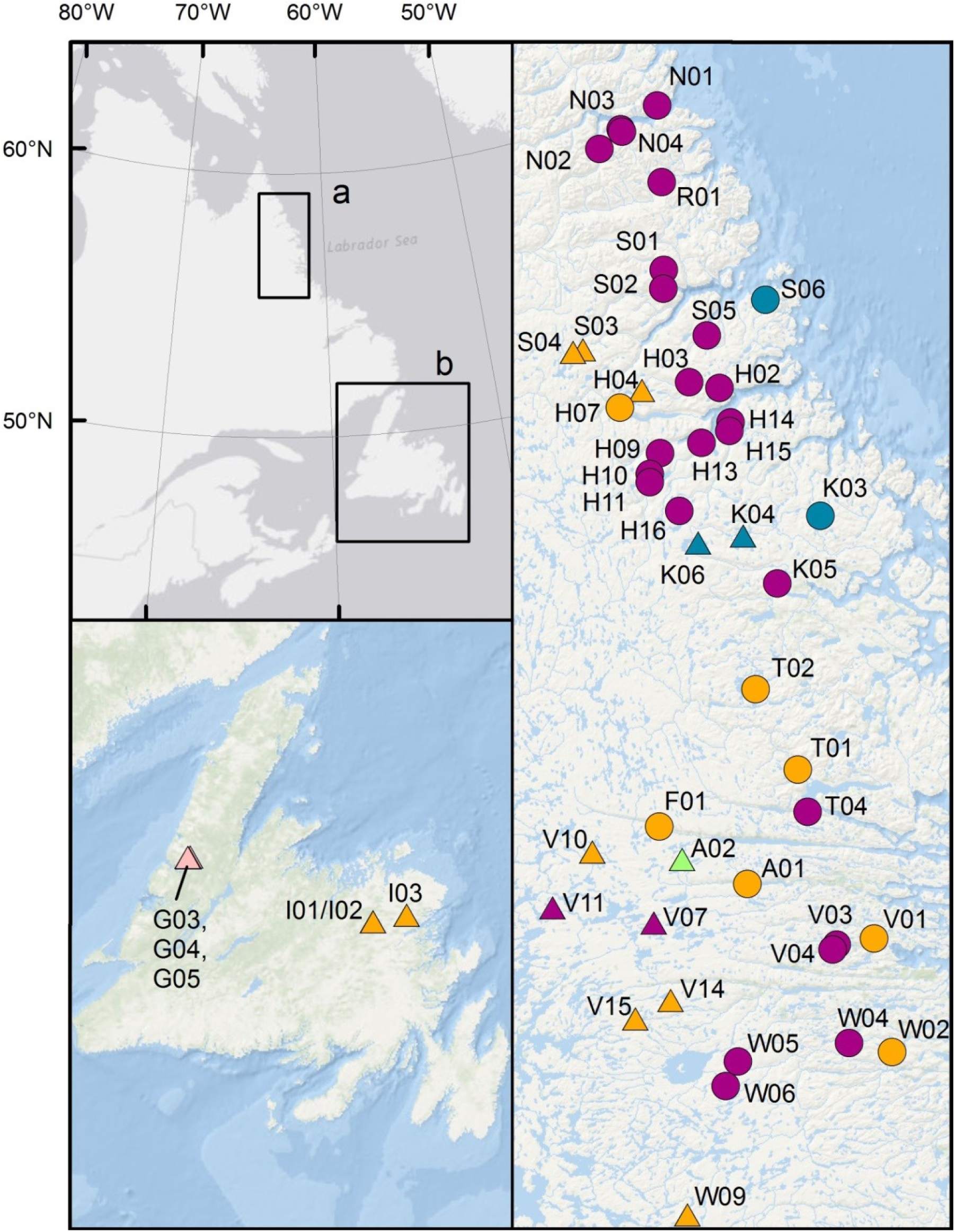
Results of SAMOVA analysis when considering all locations in a) Labrador and b) Newfoundland with > 10 samples and taking into account geography of locations using a Delaunay matrix. Locations are coloured by grouping (K = 5). Sea-accessible sites are denoted by circles, landlocked sites are denoted by triangles. Map created using ArcGIS (ESRI).

## DISCUSSION

### Extent of Secondary Contact

Our results indicate an extensive overlap in the contemporary ranges of the Arctic, Atlantic and Acadian lineages in Newfoundland and Labrador. A SAMOVA detected groupings from geographically separate locations which indicates the widespread distribution of these three lineages. The two detected groupings with the highest number of locations (orange and purple groups in Fig.4) spanned the entirety of the latitudinal range sampled in Labrador. The orange group (Fig.4), which included locations with predominately Atlantic lineage fish, also included the eastern Newfoundland locations, reflecting the extensive colonization of the Atlantic lineage throughout Newfoundland and Labrador.

The Arctic and Atlantic lineages exhibited extensive secondary contact within Labrador. Contrary to our hypothesis, there was no association between latitude and presence of the Arctic or Atlantic lineage in Labrador, suggesting that the secondary contact between these two lineages is at least as extensive as our study area. Our results represent the furthest north an Atlantic haplotype has been observed in Labrador. This observation is consistent with evidence for an incursion of Atlantic lineage nuclear DNA, but not mtDNA, in Nunavut (Moore et al. 2015). Our results also include the furthest south an Arctic lineage haplotype has been observed in the Atlantic. It is possible that the Arctic lineage may have colonized even further south into Labrador than the range considered here.

While the majority of locations in Labrador contained both the Arctic and Atlantic lineages, four populations in Labrador contained exclusively Arctic lineage samples. This group of four lakes was detected as significant by SAMOVA (blue locations in Fig.4). The absence of Atlantic lineage haplotypes (which are present in nearby populations) from these locations may be due to sampling bias or they may have been lost through drift. Alternatively, colonization by the Atlantic lineage may have been prevented by their maladaptation to these sites (“isolation-by-adaptation”) or their exclusion by the previously established Arctic lineage (“isolation-by-colonization”) (Waters 2011, Orsini et al. 2013, Waters et al. 2013).

Unlike the Atlantic lineage, the Arctic lineage does not appear to have invaded Newfoundland. The absence of the Arctic lineage from Newfoundland may be due to our low sample size. However, our study confirmed the previous observations of the Atlantic and Acadian lineage in eastern Newfoundland (Brunner et al. 2001, Moore et al. 2015). We also demonstrated that the contemporary range of both the Atlantic and Acadian lineages extends to western Newfoundland.

Our extension of the Acadian lineage’s contemporary presence into Labrador counters previous suggestions of its relatively conserved range from its putative refugium near the northeastern United States (e.g. Brunner et al. 2001, Esin and Markevich 2018). This brings the scale of the contemporary range of the Acadian lineage in line with those observed in other char lineages.

### Evidence for Introgression

Our results here suggest extensive secondary contact but also the introgression of the Arctic, Atlantic and Acadian lineages in Newfoundland and Labrador. Many sampled locations contained at least two glacial lineages suggesting the potential for hybridization among lineages. Furthermore, six sea-accessible locations in Nachvak and Saglek fjords (N01-N04, S01-S02) contained both Arctic and Atlantic lineage individuals yet no genetic structuring was found within each of these locations based on 11 microsatellite markers in Salisbury et al. (2017). This suggests that these lineages have fully introgressed.

Hybridization between these lineages may seem surprising given that Arctic lineage is thought to have split off from all other lineages between 716 000 and 1 432 000 years BP based on mtDNA (Moore et al. 2015). Alternatively, Esin and Markevich (2018) estimate the divergence of the Arctic lineage at 400 000 – 700 000 years BP during the Nebraskan-Kansan cooling. During this time the Canadian Arctic archipelago (the putative refugium for the Arctic lineage) was separated from a refugium in the Bering Sea (Esin and Markevich 2018). Many species have demonstrated reproductive isolation between different glacial lineages upon secondary contact within such a time scale (Hewitt 1996, 2003; Bernatchez and Wilson 1998, Schluter 1998). However, our results support previous research suggesting hybridization among Arctic char glacial lineages. Atlantic lineage nuclear DNA has been found in Nunavut populations of Arctic lineage individuals (Moore et al. 2015). A similar lack of a relationship between mtDNA and nuclear DNA has also been observed in three-spine stickleback (Lescak et al. 2017). Many *Salvelinus* species are known to readily hybridize (Taylor 2003), and there is evidence for Arctic char having hybridized with brook trout in Quebec (Bernatchez et al. 1995, Glémet et al. 1998) and Labrador (Fraser River) (Hammar et al. 1991) and with lake trout in Nunavut (Wilson and Hebert 1993) and Quebec (Wilson and Bernatchez 1998). These hybridizations overcome a much older allopatric divergence further supporting the introgression of the relatively recently diverged glacial lineages of Arctic char.

Some of the brook trout and lake trout mtDNA haplotypes detected in our samples may therefore reflect hybridization or backcrosses between these species and Arctic char. This would require further validation using nuclear markers but was beyond the scope of this study. An open area for future investigation is the degree to which genes from lake trout and brook trout have introgressed into Arctic char genomes within this region.

### Colonization History

We detected several rare haplotypes that were previously found in other populations within each lineage’s respective range allowing for insight into the origins of the three glacial lineages in this region. The ARC20 and ARC22 haplotypes we detected in Labrador were previously observed in geographically distinct locations across the high Canadian Arctic (Moore et al. 2015). The Arctic lineage may have therefore colonized Labrador multiple times from geographically distant populations. The ATL19 haplotype we observed in a single dark char morph in Gander Lake was previously observed in an unspecified morph in this lake as well as in a resident lacustrine population from Scotland (Moore et al. 2015). Lastly, the ATL31 haplotype we found in multiple anadromous populations was also found in a landlocked, Swedish population (Oleinik et al. 2017). The appearance of these Atlantic haplotypes on opposite sides of the Atlantic Ocean suggests extensive colonization throughout the Atlantic from the Atlantic refugium. While our study area demonstrates a high diversity of Atlantic lineage haplotypes, this diversity is no doubt due to our intensive sampling. The exact location of the Atlantic refugium remains to be determined.

### Landlocked vs. Sea-accessible locations

It was not possible to determine the order in which the glacial lineages colonized Labrador based on the lineages present in landlocked versus sea-accessible locations. All three lineages were present in both land-locked and sea-accessible locations in Labrador. Moore et al. (2015) suggested the Atlantic lineage had colonized the high Canadian Arctic after the Arctic lineage since some anadromous char populations contained Atlantic lineage nuclear DNA but nearby landlocked char populations demonstrated Arctic lineage nuclear DNA. Our results suggest that all three lineages may have colonized Labrador around the same time.

Though they did not share a common lineage, most landlocked populations contained a single lineage and low haplotypic diversity. This could be due to a founder-take-all scenario, where the lineage that first colonized a lake rapidly expanded to fill available habitat, preventing subsequent incursions from other lineages (Waters 2011, Orsini et al. 2013, Waters et al 2013). Also, landlocked populations, are more isolated and tend to exhibit smaller effective sizes (see Salisbury et al. 2017) and thus experience more drift than anadromous populations potentially leading to a greater loss of mtDNA haplotypes.

Several landlocked lakes countered this trend of reduced diversity. Landlocked lakes within the Kogaluk River system (i.e. V10, V11, V15, V16) had Arctic and Atlantic lineage char co-occurring. Access to this watershed may have been enhanced by significant run-off from the paleolake Naskaupi, which drained through the Kogaluk 7 500 and 6 000 years BP (Barnett and Peterson 1964, Jansson and Kleman 2004). Alternatively, many lakes within the Kogaluk River drainage are connected via shallow streams which could facilitate the occasional migration between lakes as it has for lake trout (McCracken et al. 2013) and longnose suckers (*Catostomus catostomus*) (Salisbury et al. 2016) in this system. Migration may have countered genetic drift (Tallmon et al. 2004), maintaining both Arctic and Atlantic lineage haplotypes in these lakes. High effective sizes (Gomez-Uchida et al. 2013) and high migration among lakes (Gomez-Uchida et al. 2009) may have similarly countered the effects of genetic drift in landlocked populations in western Newfoundland (G02-G04) which contained both Atlantic and Acadian lineages as well as high haplotypic diversity within the Acadian lineage.

### Glacial lineage and contemporary morph divergence in Gander Lake

Previous work has suggested that the high degree of neutral genetic differences observed between pale and dark morph char could be ascribed to differential glacial origins (Gomez-Uchida et al. 2008). Our results, indicating that most char in Gander Lake were of the Atlantic lineage regardless of morph, reject this hypothesis. This suggests that the great morphological, ecological and genetic differences between the pale and dark morph (O’Connell and Dempson 2002, Power et al. 2005, Gomez-Uchida et al. 2008) may have arisen in sympatry in Gander Lake within the last ∼ 10 000 years since its deglaciation (Bryson et al. 1969, Dyke 2004, Shaw et al. 2006). This is consistent with the presumed sympatric divergence of other lacustrine Arctic char morphs (Magnusson and Ferguson 1987, Volpe and Ferguson 1996, Gíslason et al. 1999). The large genetic divergence among pale and dark morph char in Gander suggests substantial genetic differences can accumulate between morphs within a short period of time, potentially fueled by divergent selection (Taylor 2003, Bernatchez 2003) and the relatively low effective population sizes of both pale and dark char (Gomez-Uchida et al. 2008). The very large genetic differences observed between these morphs (Gomez-Uchida et al. 2008) may be due to reproductive isolation resulting from a chromosomal incompatibility such as a chromosomal inversion as has been shown for other genetically, phenotypically, and ecologically differentiated fish morphs (Wellenreuther and Bernatchez (2018) and references therein).

The occurrence of Atlantic and Acadian lineages in the pale morph suggests introgression of these lineages. Similar evidence for introgression among the Arctic and Atlantic lineages was found in R01 by Salisbury et al. (2017), where morphologically-identified anadromous and resident char were found to be genetically differentiated by STRUCTURE but each contained both Arctic and Atlantic lineage individuals. Populations of sympatric dwarf and normal whitefish (*Coregonus clupeaformis*) in Maine have also each demonstrated both of two mtDNA haplotype groups (indicative of two glacial lineages) (Bernatchez and Dodson 1990, Pigeon et al. 1997). These observations lead to the puzzling implication that glacial lineages have introgressed despite thousands of years of allopatric divergence yet, in some cases, their descendants have become reproductively isolated (perhaps in sympatry) and subsequently significantly diverged in the (relatively) short time since deglaciation.

### Management Implications and the Utility of Intensive mtDNA Sampling

The likely introgression among glacial lineages in Labrador has important implications for the char fishery in Labrador. There was evidence of Arctic, Atlantic and even Acadian lineage fish in sea-accessible locations in the Notakwonan, Voisey, Anaktalik, Nain, and Okak drainages. These populations probably contribute to the commercial fishery stock complexes (Coady and Best 1976, LeDrew 1980, DFO 2001, Dempson et al. 2008). The expected introgression between lineages suggests that there is likely no need to manage them separately.

Our results verify the utility of intensive mtDNA sampling across many populations, particularly within a secondary contact zone. This approach facilitated the detection of a number of new haplotypes for the Arctic, Atlantic and Acadian lineages (Fig.2, 3) as well as the detection for the first time, of the Acadian lineage within Labrador. Finally, our detection of non-Arctic char salmonid species highlights the morphological ambiguity of salmonids, particularly as juveniles. All of the samples identified genetically as a species other than Arctic char had a median length of 40 mm (data not shown). Since species misidentification can have repercussions for the interpretation of genetic data we therefore caution against the exclusive use of morphology in juveniles in regions where other salmonids coexist with Arctic char. The mtDNA-based technique as used here is useful for minimizing the possibility of species misidentification in regions where other salmonid species overlap with Arctic char.

In conclusion, our results clearly demonstrate the widespread secondary contact of the Arctic, Atlantic, and Acadian glacial lineages of Arctic char throughout Newfoundland and Labrador, Canada. These three glacial lineages have likely introgressed extensively in this region. The genetic divergence in morph pairs in Ramah and Gander lakes do not appear to be linked to glacial lineages. We demonstrate that Arctic char are an ideal model species for future investigation of secondary contact zones and the influence of historical allopatry on contemporary genetic structure and niche divergence.

## ACKNOWLEDGEMENTS

Thanks go to S. Avery, H. Buchanan, D. Cote, J. Callahan, T. Gallant, S. Gerrow, B. Green, S. Hann, T. Hann, L. House, T. Knight, J. Merkuratsuk, F. Palstra, L. Pike, R. Reid, J. Seibert, A. Simpson, R. Solomon, D. Gomez-Uchida, A. Walsh, C. Webb, and J. Webb for their indispensable help with field work. Thanks also to D. Notte for her help with lab work. We greatly appreciate Parks Canada for allowing us access to the Torngat Mountains National Park and the Nunatsiavut government for allowing us to access their lands. We also thank the Institute for Biodiversity, Ecosystem Science and Sustainability of the Department of Environment and Conservation of the Government of Newfoundland and Labrador for funding for this project; NSERC for the Strategic Grant STPGP 430198 and Discovery Grant awarded to DER, for the CGS-D awarded to SJS; the Killam Trust for the Level 2 Izaak Walton Killam Predoctoral Scholarship awarded to SJS; and the Government of Nova Scotia for the Graduate Scholarship awarded to SJS.

## REFERENCES

Anderson, T.C. 1985. Rivers of Labrador. Can. Spec. Publ. Fish. Aquat. Sci. 81, Ottawa. 389 pp.

Bandelt, H. J., Forster, P., and Röhl, A. 1999. Median-joining networks for inferring intraspecific phylogenies. Mol. Biol. Evolution. 16(1), 37–48. doi: https://doi.org/10.1093/oxfordjournals.molbev.a026036.

Barnett, D.M., and Peterson, J.A. 1964. The Significance of Glacial Lake Naskaupi in the Deglaciation of Labrador-Ungava. Can. Geogr. 8(4), 173–181. doi: https://doi.org/10.1111/j.1541-0064.1964.tb00606.x.

Bernatchez, L. 2003. Ecological Theory of Adaptive Radiation: An Empirical Assessment from Coregonine Fishes (Salmonifirnes). Evolution illuminated: salmon and their relatives (eds Stearns, S.C. and Hendry, A.P.), pp. 176–207. Oxford University Press, Oxford.

Bernatchez, L., Dempson, J.B., and Martin, S. 1998. Microsatellite gene diversity analysis in anadromous arctic char, *Salvelinus alpinus*, from Labrador, Canada. Can. J. Fish. Aquat. Sci. 55(5), 1264–1272. doi: https://doi.org/10.1139/f97-325.

Bernatchez, L., and Dodson, J.J. 1990. Allopatric origin of sympatric populations of lake whitefish (*Coregonus clupeaformis*) as revealed by mitochondrial-DNA restriction analysis. Evolution. 44(5), 1263–1271. doi: https://doi.org/10.1111/j.1558-5646.1990.tb05230.x.

Bernatchez, L., Glémet, H., Wilson, C.C., and Danzmann, R.G. 1995. Introgression and fixation of Arctic char (*Salvelinus alpinus*) mitochondrial genome in an allopatric population of brook trout (*Salvelinus fontinalis*). Can. J. Fish. Aquat. Sci. 52(1), 179–185. doi: https://doi.org/10.1139/f95-018.

Bernatchez, L., and Wilson, C.C. 1998. Comparative phylogeography of Nearctic and Palearctic fishes. Mol. Ecol. 7(4), 431–452. doi: https://doi.org/10.1046/j.1365-294x.1998.00319.x.

Brown Gladden, J.G., Postma Maiers, L.D., Carmichael, T.J., and Reist, J.D. 1995. Mitochondrial DNA Control Region Sequence Variation in Arctic Char (*Salvelinus alpinus* (L.)). Canadian Data Report of Fisheries and Aquatic Sciences 968. Department of Fisheries and Oceans, Central and Arctic Region.

Brunner, P.C., Douglas, M.R., Osinov, A., Wilson, C.C., and Bernatchez, L. 2001. Holarctic phylogeography of Arctic charr (*Salvelinus alpinus* L.) inferred from mitochondrial DNA sequences. Evolution. 55(3), 573–586. doi: https://doi.org/10.1554/0014-3820(2001)055[0573:HPOACS]2.0.CO;2.

Bryson, R.A., Wendland, W.M., Ives, J.D., and Andrews, J.T. 1969. Radiocarbon Isochrones on the Disintegration of the Laurentide Ice Sheet. Arctic Alpine Res. 1(1), 1–13. doi: 10.1080/00040851.1969.12003535.

Coady L.W., and Best C.W. 1976. Biological and management investigations of the Arctic char fishery at Nain, Labrador. Can. Fish Mar. Serv. Res. Dev. Tech. Rep. 624, 103 pp.

Delaunay, B. 1934. Sur la sphère vide. B. Acad. Sci USSR, 7, 793–800.

DFO. 2001. North Labrador Arctic Charr. DFO Science Stock Status Report D2–07.

Dupanloup, I., Schneider, S., and Excoffier, L. 2002. A simulated annealing approach to define the genetic structure of populations. Mol. Ecol. 11(12), 2571–2581. doi: https://doi.org/10.1046/j.1365-294X.2002.01650.x.

Dyke, A.S. 2004. An outline of North American deglaciation with emphasis on central and northern Canada. In Developments in Quaternary Sciences (eds Ehlers, J., Gibbard, P. L.), Vol. 2, pp. 373–424. Elsevier, Amsterdam.

Esin, E.V., Bocharova, E.S., Mugue, N.S., and Markevich, G.N. 2017. Occurrence of sympatric charr groups, *Salvelinus*, Salmonidae, in the lakes of Kamchatka: a legacy of the last glaciations. J. Fish Biol. 91(2), 628–644. doi: 10.1111/jfb.13378.

Esin, E.V., and Markevich, G.N. 2018. Evolution of the Charrs, Genus *Salvelinus* (Salmonidae). Origins and Expansion of the Species. J. Ichthyol. 58(2), 187–203. doi: https://doi.org/10.1134/S0032945218020054.

Fraser C.I., Nikula, R., Ruzzante, D.E., and Waters, J.M. 2012. Poleward bound: biological impacts of Southern Hemisphere glaciation. Trends in Ecology and Evolution, August 2012, Vol. 27, No. 8: 462 – 471. doi: http://dx.doi.org/10.1016/j.tree.2012.04.011.

Gíslason, D., Ferguson, M.M., Skúlason, S., and Snorrason, S.S. 1999. Rapid and coupled phenotypic and genetic divergence in Icelandic Arctic char (*Salvelinus alpinus*). Can. J. Fish. Aquat. Sci. 56(12), 2229–2234. doi: https://doi.org/10.1139/f99-245.

Glémet, H., Blier, P., and Bernatchez, L. 1998. Geographical extent of Arctic char (*Salvelinus alpinus*) mtDNA introgression in brook char populations (*S. fontinalis*) from eastern Quebec, Canada. Mol. Ecol. 7(12), 1655–1662. doi: https://doi.org/10.1046/j.1365-294x.1998.00494.x.

Gomez-Uchida, D., Dunphy, K.P., O’connell, M.F., and Ruzzante, D.E. 2008. Genetic divergence between sympatric Arctic charr *Salvelinus alpinus* morphs in Gander Lake, Newfoundland: roles of migration, mutation and unequal effective population sizes. J. Fish Biol. 73(8), 2040–2057. doi: 10.1111/j.1095-8649.2008.02048.x.

Gomez-Uchida, D., Knight, T.W., and Ruzzante, D.E. 2009. Interaction of landscape and life history attributes on genetic diversity, neutral divergence and gene flow in a pristine community of salmonids. Mol. Ecol. 18(23), 4854–4869. doi: 10.1111/j.1365-294X.2009.04409.x.

Gomez-Uchida, D., Palstra, F. P., Knight, T.W., and Ruzzante, D.E. 2013. Contemporary effective population and metapopulation size (Ne and meta-Ne): comparison among three salmonids inhabiting a fragmented system and differing in gene flow and its asymmetries. Ecol. Evol. 3(3), 569–580. doi: 10.1002/ece3.485.

Guindon, S., and Gascuel, O. 2003. A simple, fast, and accurate algorithm to estimate large phylogenies by maximum likelihood. Syst. Biol. 52(5), 696–704. doi:10.1080/10635150390235520.

Hammar, J., and Filipsson, O. 1985. Ecological test fishing with the Lundgren gillnets of multiple mesh size: the Drottningholm technique modified for Newfoundland Arctic char populations. Report, Institute of Freshwater Research, Drottningholm. 62, 12–35.

Hammar, J., Dempson, J.B., and Verspoor, E. 1991. Natural hybridization between Arctic char (*Salvelinus alpinus*) and brook trout (*S. fontinalis*): evidence from northern Labrador. Can. J. Fish. Aquat. Sci. 48(8), 1437–1445. doi: https://doi.org/10.1139/f91-171.

Hewitt, G.M. 1988. Hybrid zones-natural laboratories for evolutionary studies. Trends Ecol. Evol. 3(7), 158–167. doi: 10.1016/0169-5347(88)90033-X.

Hewitt, G.M. 1996. Some genetic consequences of ice ages, and their role in divergence and speciation. Biol. J. Linn. Soc. 58(3), 247–276. doi: https://doi.org/10.1111/j.1095-8312.1996.tb01434.x.

Hewitt, G. 2000. The genetic legacy of the Quaternary ice ages. Nature. 405(6789), 907–913. doi: https://doi.org/10.1038/35016000.

Hewitt, G. 2003. Ice ages: species distributions, and evolution. In Evolution on planet Earth (eds Rothschild, L. J., Lister A. M.), pp. 339–361. Academic Press, London. doi: https://doi.org/10.1016/B978-012598655-7/50045-8.

Hewitt, G.M. 2004. Genetic consequences of climatic oscillations in the Quaternary. Philos. T. R. Soc. B. 359(1442), 183–195. doi: 10.1098/rstb.2003.1388.

Jansson, K.N. 2003. Early Holocene glacial lakes and ice marginal retreat pattern in Labrador/Ungava, Canada. Palaeogeogr. Palaeocl. 193(3-4), 473–501. doi: https://doi.org/10.1016/S0031-0182(03)00262-1.

Jansson, K.N., and Kleman, J. 2004. Early Holocene glacial lake meltwater injections into the Labrador Sea and Ungava Bay. Paleoceanography. 19(1), PA1001. doi: https://doi.org/10.1029/2003PA000943.

LeDrew, L.J. 1980. Research on the Arctic char fishery of northern Labrador. Can.MS Rep. Fish. Aquat. Sci. 1589.

Leigh, J.W., and Bryant, D. 2015. Popart: full-feature software for haplotype network construction. Methods Ecol. Evol. 6(9), 1110–1116. doi: https://doi.org/10.1111/2041-210X.12410.

Lescak, E.A., Wund, M.A., Bassham, S., Catchen, J., Prince, D.J., Lucas, R., Dominguez, G., von Hippel, F.A., and Cresko, W.A. 2017. Ancient three-spined stickleback (*Gasterosteus aculeatus*) mtDNA lineages are not associated with phenotypic or nuclear genetic variation. Biol. J. Linn. Soc. 122(3), 579–588. doi: https://doi.org/10.1093/biolinnean/blx080.

Magnusson, K.P., and Ferguson, M.M. 1987. Genetic analysis of four sympatric morphs of Arctic charr, *Salvelinus alpinus*, from Thingvallavatn, Iceland. Environ. Biol. Fish. 20(1), 67–73. doi: https://doi.org/10.1007/BF00002026.

McCracken, G.R., Perry, R., Keefe, D., and Ruzzante, D.E. 2013. Hierarchical population structure and genetic diversity of lake trout (*Salvelinus namaycush*) in a dendritic system in Northern Labrador. Freshwater Biol. 58(9), 1903–1917. doi: https://doi.org/10.1111/fwb.12179.

Moore, J.S., Bajno, R., Reist, J.D., and Taylor, E.B. 2015. Post-glacial recolonization of the North American Arctic by Arctic char (*Salvelinus alpinus*): genetic evidence of multiple northern refugia and hybridization between glacial lineages. J. Biogeogr. 42(11), 2089–2100. doi: https://doi.org/10.1111/jbi.12600.

Noakes, D.L. 2008. Charr truth: sympatric differentiation in *Salvelinus* species. Environ. Biol. of Fish. 83(1), 7–15. doi: https://doi.org/10.1007/s10641-008-9379-x.

Noor, M.A. 1999. Reinforcement and other consequences of sympatry. Heredity. 83(5), 503–508. doi: 10.1046/j.1365-2540.1999.00632.x.

Occhietti, S., Parent, M., Lajeunesse, P., Robert, F., and Govare, É. 2011. Late Pleistocene–Early Holocene decay of the Laurentide Ice Sheet in Québec–Labrador. In Developments in Quaternary Sciences, Vol. 15, pp. 601–630. Elsevier. doi: https://doi.org/10.1016/B978-0-444-53447-7.00047-7.

O’Connell, M.F., and Dempson, J.B. 2002. The biology of Arctic charr, Salvelinus alpinus, of Gander Lake, a large, deep, oligotrophic lake in Newfoundland, Canada. In Ecology, behaviour and conservation of the charrs, genus Salvelinus (eds Magnan, P., Audet, C., Glémet, H., Rodriguez, M.A., Taylor, E.B.), pp. 115–126. Springer, Dordrecht. doi: https://doi.org/10.1023/A:1016001423937.

Oleinik, A.G., Skurikhina, L.A., and Kukhlevsky, A.D. 2017. Secondary contact between two divergent lineages of charrs of the genus *Salvelinus* in the Northwest Pacific. Russ. J. Gen+. 53(11), 1221–1233. doi: https://doi.org/10.1134/S1022795417110084.

Orsini, L., Vanoverbeke, J., Swillen, I., Mergeay, J., and Meester, L. 2013. Drivers of population genetic differentiation in the wild: isolation by dispersal limitation, isolation by adaptation and isolation by colonization. Mol. Ecol. 22(24), 5983–5999. doi: https://doi.org/10.1111/mec.12561.

Pigeon, D., Chouinard, A., and Bernatchez, L. 1997. Multiple modes of speciation involved in the parallel evolution of sympatric morphotypes of lake whitefish (*Coregonus clupeaformis*, Salmonidae). Evolution. 51(1), 196–205. doi: https://doi.org/10.1111/j.1558-5646.1997.tb02401.x.

Piry, S., Alapetite, A., Cornuet, J.M., Paetkau, D., Baudouin, L., and Estoup, A. 2004. GENECLASS2: a software for genetic assignment and first-generation migrant detection. J. Hered. 95(6), 536–539. doi: https://doi.org/10.1093/jhered/esh074.

Power, G. 2002. Charrs, glaciations and seasonal ice. In Ecology, behaviour and conservation of the charrs, genus Salvelinus (pp. 17–35). Springer, Dordrecht. doi: https://doi.org/10.1007/978-94-017-1352-8_2.

Power, M., O’Connell, M.F., and Dempson, J.B. 2005. Ecological segregation within and among Arctic char morphotypes in Gander Lake, Newfoundland. Environ. Biol. Fish. 73(3), 263–274. doi: https://doi.org/10.1007/s10641-005-2137-4.

R Development Core Team. 2013. R: a language and environment for statistical computing. R Foundation for Statistical Computing, Vienna, Austria.

D. Ruzzante, D.E., Walde, S.J., Gosse, J.C., Cussac, V.E., Habit, E., Zemlak, T.S., and Adams, E. 2008. Climate control on ancestral population dynamics: insight from Patagonian fish phylogeography. Mol. Ecol. 17(9), 2234–2244. doi: 10.1111/j.1365-294X.2008.03738.x.

Salisbury, S.J., McCracken, G.R., Keefe, D., Perry, R., and Ruzzante, D.E. 2016. A portrait of a sucker using landscape genetics: how colonization and life history undermine the idealized dendritic metapopulation. Mol. Ecol. 25(17), 4126–4145. doi: https://doi.org/10.1111/mec.13757.

E. Salisbury, S.J., Booker, C., McCracken, G.R., Knight, T., Keefe, D., Perry, R., and Ruzzante, D. 2017. Genetic divergence among and within Arctic char (*Salvelinus alpinus*) populations inhabiting landlocked and sea-accessible sites in Labrador, Canada. Can. J. Fish. Aquat. Sci. 999, 1–14. doi: https://doi.org/10.1139/cjfas-2017-0163.

Schluter D. 1998. Ecological causes of speciation. In: Endless Forms: Species and Speciation (eds Howard, D. J., Berlocher, S. H.), pp. 114–129. Oxford University Press, New York. doi: https://doi.org/10.1086/285901.

Schluter, D. 2001. Ecology and the origin of species. Trends Ecol. Evol. 16(7), 372–380. doi: https://doi.org/10.1016/S0169-5347(01)02198-X.

Scott, W.B., and Crossman, E.J. 1973. Freshwater fishes of Canada. Bull. Fish. Res. Board Can. No. 184. Galt House Publication Ltd., Oakville. 966 pp.

Shaw, J., Piper, D.J.W., Fader, G.B.J., King, E.L., Todd, B.J., Bell, T., Batterson, M.J., and Liverman, D.G.E. 2006. A conceptual model of the deglaciation of Atlantic Canada. Quaternary Sci. Rev. 25(17-18), 2059–2081. doi: https://doi.org/10.1016/j.quascirev.2006.03.002.

Soltis, D.E., Morris, A.B., McLachlan, J.S., Manos, P.S., and Soltis, P.S. 2006. Comparative phylogeography of unglaciated eastern North America. Mol. Ecol. 15(14), 4261–4293. doi: https://doi.org/10.1111/j.1365-294X.2006.03061.x.

Swenson, N.G., and Howard, D.J. 2005. Clustering of contact zones, hybrid zones, and phylogeographic breaks in North America. Am. Nat. 166(5), 581–591. doi: https://doi.org/10.1086/491688.

Tallmon, D.A., Luikart, G., and Waples, R.S. 2004. The alluring simplicity and complex reality of genetic rescue. Trends Ecol. Evol. 19(9), 489–496. doi: https://doi.org/10.1016/j.tree.2004.07.003.

Taylor, E.B. 2003. Evolution in mixed company: evolutionary inferences from studies of natural hybridization in Salmonidae. In Evolution illuminated: salmon and their relatives (eds Stearns, S.C. and Hendry, A. P.), pp. 232–263. Oxford University Press, Oxford.

Volpe, J.P., and Ferguson, M.M. 1996. Molecular genetic examination of the polymorphic Arctic charr *Salvelinus alpinus* of Thingvallavatn, Iceland. Mol. Ecol. 5(6), 763–772. doi: https://doi.org/10.1111/j.1365-294X.1996.tb00372.x.

Waters J.M. 2011. Competitive exclusion: phylogeography’s ‘elephant in the room’? Mol. Ecol. 20, 4388–4394. doi: https://doi.org/10.1111/j.1365-294X.2011.05286.x.

Waters, J.M., Fraser, C.I., and Hewitt, G.M. 2013. Founder takes all: density-dependent processes structure biodiversity. Trends Ecol. Evol. 28(2): 78–85. doi: https://doi.org/10.1016/j.tree.2012.08.024.

Wellenreuther, M., and Bernatchez, L. 2018. Eco-evolutionary genomics of chromosomal inversions. Trends Ecol. Evol. 33(6): 427–440. doi: https://doi.org/10.1016/j.tree.2018.04.002.

Wilson, C.C., and Bernatchez, L. 1998. The ghost of hybrids past: fixation of arctic charr (*Salvelinus alpinus*) mitochondrial DNA in an introgressed population of lake trout (*S. namaycush*). Mol. Ecol. 7(1), 127–132. doi: https://doi.org/10.1046/j.1365-294x.1998.00302.x.

Wilson, C.C., and Hebert, P.D. 1993. Natural hybridization between Arctic char (*Salvelinus alpinus*) and lake trout (*S. namaycush*) in the Canadian Arctic. Can. J. Fish. Aquat. Sci. 50(12), 2652–2658. doi: https://doi.org/10.1139/f93-288.

Wilson, C.C., Hebert, P.D.N., Reist, J.D., and Dempson, J. 1996. Phylogeography and postglacial dispersal of arctic charr *Salvelinus alpinus* in North America. Mol. Ecol. 5(2), 187–197. doi: https://doi.org/10.1046/j.1365-294X.1996.00265.x.

